# Genetic variation, environment and demography intersect to shape Arabidopsis defense metabolite variation across Europe

**DOI:** 10.1101/2020.09.23.310268

**Authors:** Ella Katz, Clement Bagaza, Samuel Holden, Ruthie Angelovici, Daniel J. Kliebenstein

## Abstract

Plants face a variety of challenges within their ever-changing environment. Diverse metabolites are central to the plants ability to overcome these challenges. Understanding the environmental and genetic factors influencing the variation in specialized metabolites is the key to understand how plants survive and develop under changing environments. Here we measure the variation in specialized metabolites across a population of 797 natural *Arabidopsis thaliana* accessions. We show a combination of geography, environmental parameters, demography, and different genetic processes that creates a specific pattern in their accumulation and distribution. By identifying and tracking causal polymorphisms at multiple loci controlling metabolites variation we show that each locus displays extensive allelic heterogeneity with signatures of both parallel and convergent evolutionary processes. These loci combine epistatically and show differing relationships to environmental parameters leading to different distributions. This provides a detailed perspective about the complexity of the forces and mechanisms that shape the accumulation and distribution of a family of specialized metabolites critical for plant fitness.

## Introduction

The biotic and abiotic components of a plant’s habitat/environment are continuously changing. This creates a complex system to which a plant must develop adaptation strategies to ensure survival and reproduction. Metabolites are frequent keys to these strategies, involving the production and accumulation of different metabolites from signaling hormones, primary metabolites and a wide array of multi-functional specialized metabolites (Erb & Kliebenstein, 2020; Hanower & Brzozowska, 1975; Hayat et al., 2012; Kim et al., 2012; D J Kliebenstein, 2004; Malcolm, 1994; Thakur & Rai, 1982; Wolters & Jürgens, 2009; Yang, Lin, & Kao, 2000). The complete suite of these metabolites helps to determine the plants survival and development. A complication in the plants ability to create an optimal blend of metabolite-based strategies is the fact that individual specialized metabolites can have contrasting effects in a complex environment. For example, individual specialized metabolites can provide defense against some attackers while simultaneously causing sensitivity to other biotic attackers or abiotic stresses (Agrawal, 2000; Bialy, Oleszek, Lewis, & Fenwick, 1990; Erb & Kliebenstein, 2020; Futuyma & Agrawal, 2009; Hu et al., 2018; Lankau, 2007; Opitz & Müller, 2009; Uremis, Arslan, Sangun, Uygur, & Isler, 2009; Züst & Agrawal, 2017). This creates offsetting ecological benefits and costs for individual metabolites that when summed across all the metabolites means that there are complex selective pressures driving the differentiation of metabolic profiles within a species and shaping genetic variation within and between populations depending on the diverse challenges faced (Fan, Leong, & Last, 2019; R. Kerwin et al., 2015; Malcolm, 1994; Sønderby, Geu-Flores, & Halkier, 2010; Szakiel, Pączkowski, & Henry, 2011; Wentzell & Kliebenstein, 2008; Züst et al., 2012).

Recent decades have seen significant advances in the identification of the genetic variation creating this metabolic variation. A common theme developing from these studies is that the metabolic variation within and between species is the result of structural variation at the enzymes responsible for the chemical structures (Chan, Rowe, Corwin, Joseph, & Kliebenstein, 2011; Chan, Rowe, & Kliebenstein, 2010; Fan et al., 2019; Kroymann, Donnerhacke, Schnabelrauch, & Mitchell-Olds, 2003; Moore et al., 2019; Schilmiller, Pichersky, & Last, 2012). These structural variants and the resulting chemical variation strongly influence plant fitness in response to a broad range of biotic interactions including at least herbivores, but also other plant species and other members of the same plant species (Bednarek & Osbourn, 2009; Brachi et al., 2015; R. E. Kerwin et al., 2017; R. Kerwin et al., 2015; Lankau & Kliebenstein, 2009; Lankau & Strauss, 2007, 2008; Lankau, 2007). Most mechanistic studies of natural variation in specialized metabolism have focused on apparent biallelic phenotypic variation linked to loss-of-function variants. However, it is not clear if biallelic genetic causation is true when extended to a large collection of individuals from wide-ranging populations within a species. If selective pressures are sufficiently strong and non-linear, it is possible to have repeated and independent generation of structural variants creating the same metabolic variation. This raises the possibility for chemical variation within a species to show hallmarks of parallel evolution, wherein phenotypically similar variants independently arise from the same genetic background. Equally it may be possible to find within-species convergent evolution, where different allele with identical metabolic consequences arise from independent genetic backgrounds through different mechanisms. Because these genetic processes are occurring simultaneously with neutral demographic processes like migration, there is a need to better understand how the intersection of environmental pressure, demography and genomic complexity gives rise to the pattern of metabolic variation across a plant species.

To better understand how genomic variation, demography and environmental pressures shape the variation of specialized metabolism within a species, we used the model Glucosinolates (GSLs) pathway. GSLs are a diverse class of specialized metabolites produced in the order Brassicales, including the model plant Arabidopsis (*Arabidopsis thaliana*), that show extensive variation between and within species across the order (Bakker, Traw, Toomajian, Kreitman, & Bergelson, 2008; Benderoth et al., 2006; Brachi et al., 2015; Chan et al., 2010; Daxenbichler et al., 1991; Halkier & Gershenzon, 2006; R. Kerwin et al., 2015; D J Kliebenstein, Gershenzon, & Mitchell-Olds, 2001; D J Kliebenstein, Kroymann, et al., 2001; D J Kliebenstein, Lambrix, Reichelt, Gershenzon, & Mitchell-Olds, 2001; James E. Rodman, Kruckeberg, & Al-Shehbaz, 1981; James Eric Rodman, 1980; Sønderby et al., 2010; Wright, Lauga, & Charlesworth, 2002). GSLs consist of a common core structure with a highly diverse side chain that determines the GSLs biological activity in defense, growth, development and abiotic stress resistance (Beekwilder et al., 2008; Hansen et al., 2008; Hasegawa, Yamada, Kosemura, Yamamura, & Hasegawa, 2000; Katz et al., 2020; Katz, Nisani, Sela, Behar, & Chamovitz, 2015; Malinovsky et al., 2017; Salehin et al., 2019; Yamada et al., 2003). The Arabidopsis-GSL system is an optimal model to study the species wide processes driving specialized metabolite variation because the identity of the whole biosynthetic pathway is known, including the major causal loci for natural variation (Benderoth et al., 2006; Brachi et al., 2015; Chan et al., 2011, 2010; Hansen, Kliebenstein, & Halkier, 2007; D J Kliebenstein, Gershenzon, et al., 2001; D. Kliebenstein, Pedersen, Barker, & Mitchell-Olds, 2002; Daniel J Kliebenstein, Figuth, & Mitchell-Olds, 2002; Kroymann & Mitchell-Olds, 2005; Pfalz, Vogel, Mitchell-Olds, & Kroymann, 2007; Sønderby et al., 2010; Wentzell et al., 2007). These major loci, have been proven to influence Arabidopsis fitness and can be linked to herbivore pressure (Brachi et al., 2015; Hansen et al., 2008; Jander, Cui, Nhan, Pierce, & Ausubel, 2001; R. E.Kerwin et al., 2017; R. Kerwin et al., 2015; Züst et al., 2012). Beyond the major causal loci, there is also evidence from genome wide association studies for highly polygenic variation in the genetic background that further contributes to modulating GSL variation (Chan et al., 2011). The public availability of over 1000 widely distributed accessions with genomic sequences provides the ability to phenotype GSL variation across a large spatial scale and query the distribution and relationship of causal haplotypes at the major GSL causal loci.

In Arabidopsis and other Brassicas, the main GSLs are Methionine-derived, Aliphatic, GSLs. Genetic variation in Aliphatic GSLs structure is controlled by natural variation at three loci, GS-Elong, GS-AOP and GS-OH with these three loci combining to create a dominant Aliphatic GSL chemotype. In addition to these expressed loci, there is a large suite of loci that can modify these dominant patterns (Brachi et al., 2015; Chan et al., 2011, 2010). GS-Elong differentially elongates the Methionine side chain by structural variation influencing the expression of divergent methylthioalkylmalate synthase enzymes (MAM) that add carbons to the side chain (Abrahams, Pires, & Schranz, 2020). In Arabidopsis, MAM2 catalyzes the addition of two carbons to the side chain, creating GSLs with 3 carbon side chains. MAM1 catalyzes the addition of three carbons to make GSLs with 4 carbon side chains (Figure 1). MAM3 (also known as MAM-L) catalyzes the addition of up to 6 carbons (D J Kliebenstein, Lambrix, et al., 2001; Kroymann et al., 2003; Mithen, Clarke, Lister, & Dean, 1995). The core pathway leads to the creation of the methylthio GSL (MT). Then, the MT will be converted to a methylsulfinyl (MSO) with a matching number of carbons (Giamoustaris & Mithen, 1996; Hansen et al., 2007). Structural variation at the GS-AOP locus leads to differential modification of the MSO by differential expression of a family of 2-oxoacid-dependent dioxygenases (2ODD). The AOP2 enzyme removes the MSO moiety leaving an alkenyl sidechain, while AOP3 leaves a hydroxyl moiety. Previous work has suggested three alleles of GS-AOP: AOP3 expressing, AOP2 expressing and a null allele (i.e. Col-0 and similar accessions) with nonfunctional copies of AOP2 and AOP3 leading to MSO accumulation, the AOP substrate (Figure 1) (Chan et al., 2010; D J Kliebenstein, Kroymann, et al., 2001; D J Kliebenstein, Lambrix, et al., 2001; Mithen et al., 1995). The 4C alkenyl side-chain can be further modified by adding a hydroxyl group at the 2C via the GS-OH 2-ODD (Figure 1) (Hansen et al., 2008). In spite of the evolutionary distance, independent variation at the same three loci influence the structural diversity in Aliphatic-GSLs within Brassica, Streptanthus and Arabidopsis (D J Kliebenstein & Cacho, 2016; Lankau & Kliebenstein, 2009). For example the C3 MAM in Arabidopsis and Brassica represent two independent lineages as are the MAMs responsible for C4 GSLs, in fact the MAM locus contains at least three independent lineages that recreate the same length variation (Abrahams et al., 2020). This indicates repeated evolution across species, but it is not clear how frequently these loci are changing within a single species or how ecological or demographic processes may shape within-species variation at these loci.

**Figure 1:**
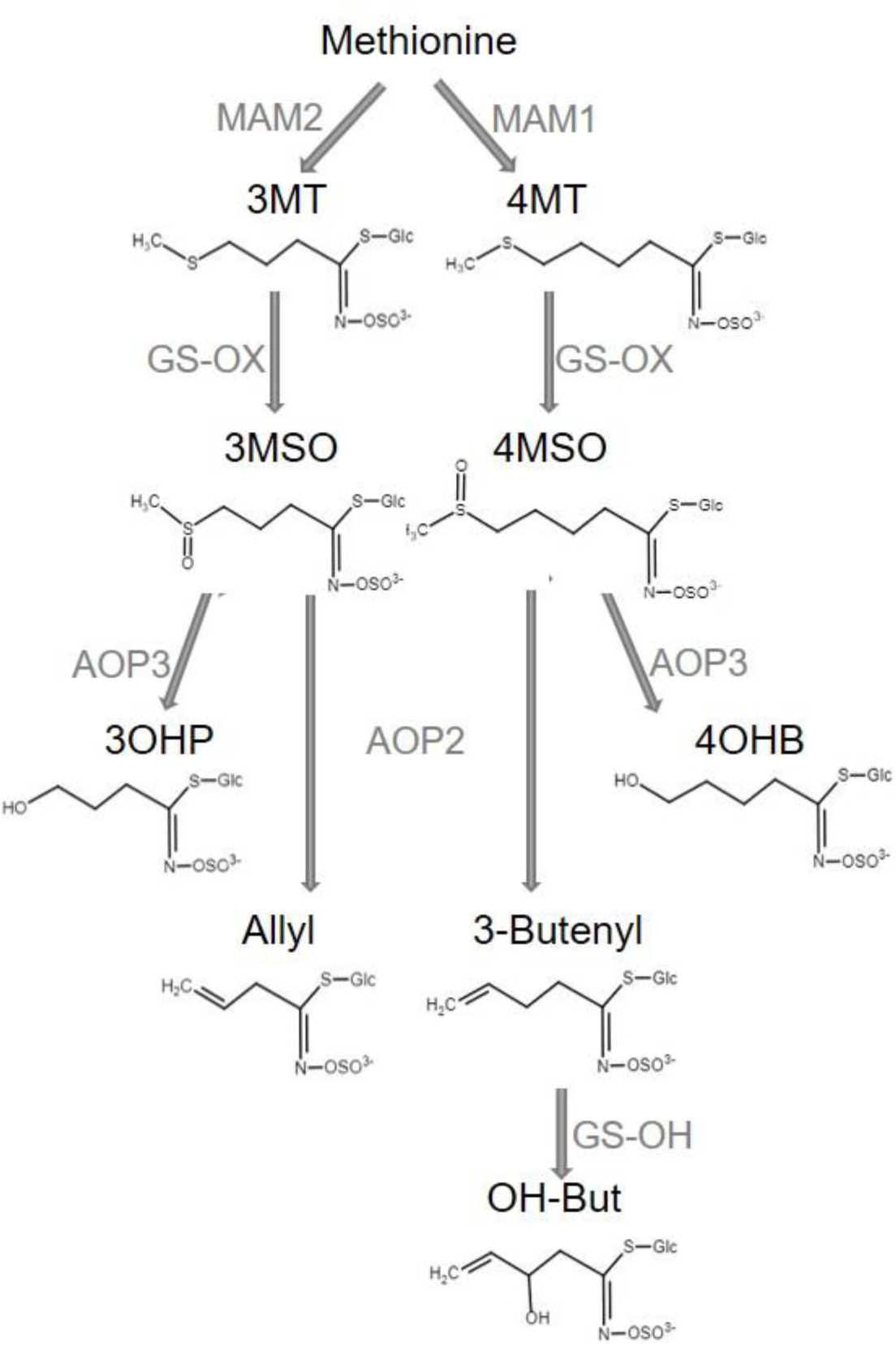
Aliphatic GSL biosynthesis pathway. Short names and structures of the GSLs are in black. Genes encoding the causal enzyme for each reaction (Arrow) are in grey. GS-OX is a gene family of five or more genes. OH-But= 2-OH-3-Butenyl.

In this work we described GSL variation in seeds of a collection of 797 *Arabidopsis thaliana* natural accessions collected from locations across Europe. The amounts of GSLs can vary across different tissues and life stages, but there is a strong correlation in the type of Aliphatic GSL produced across tissues because the major causal effect loci are not plastic due to structural variation (Brown, Tokuhisa, Reichelt, & Gershenzon, 2003; D J Kliebenstein, Gershenzon, et al., 2001; D J Kliebenstein, Kroymann, et al., 2001; Petersen, Chen, Hansen, Olsen, & Halkier, 2002). Thus, the seeds chemotype is the same as the leaves. Further, seeds have the highest level of GSLs in Arabidopsis and they are stable at room temperature until germination, hence making seeds a perfect tissue to survey variation. Further, GSLs are known to be important for seed defenses against herbivores and pathogens (Raybould & Moyes, 2001). By measuring GSLs in seeds, we identified that the distribution of GSLs and the causal alleles are influenced by a diverse set of factors with a primary contribution by geography, environmental parameters and their interaction. We describe here the complex genetic architecture of the three main causal loci responsible for the GSL composition, show how it effects the actual phenotype, and how it evolved. Interestingly, the combination of these elements reveal that several evolutional mechanisms are involved in shaping the GSL variation and distribution, and the integration of them result in the pattern described here.

## Results

### GSL variation across Europe

To investigate the genetic, environmental and demographic parameters influencing the distribution of Arabidopsis GSL chemotypes, we measured GSLs from seeds of a collection of 797 *Arabidopsis thaliana* natural accessions (The 1001 Genomes Consortium, 2016). These Arabidopsis accessions were collected from different geographical locations, mainly in and around Europe. 23 different GSLs were detected and quantified identifying a wide diversity in composition and amount among the natural accessions with a median heritability of 83%, ranging from 34% to 93% (Supplemental Table 1). To summarize the GSL variation among the accessions we performed principal component analyses (PCA) on the accumulation of all the individual GSLs across the accessions as an unbiased first step. The first two PCs only captured 33% of the total variation with PC1 describing GSLs with 4 and 7 carbons and PC2 mainly capturing GSLs with 8 carbons in their side chain (supp. Figure 1). Previous work using a collection of predominantly central European accessions had suggested a simple continental gradient chain-elongation variation from the south-west to the north-east (Brachi et al., 2015; Züst et al., 2012). To assess if this was still apparent in this larger collection, we plotted the accessions based on their geographical locations, and colored them based on their PC1 and PC2 scores that are linked to chain elongation variation (Figure 2A and supp. Figure 2A, respectively). This larger collection shows that there is not a single gradient shaping GSL diversity across Europe (Figure 2A). Instead the extended sampling of accessions around the Mediterranean in this collection shows that the SW to NE pattern reiterates within the Iberian Peninsula.

**Table 1:**
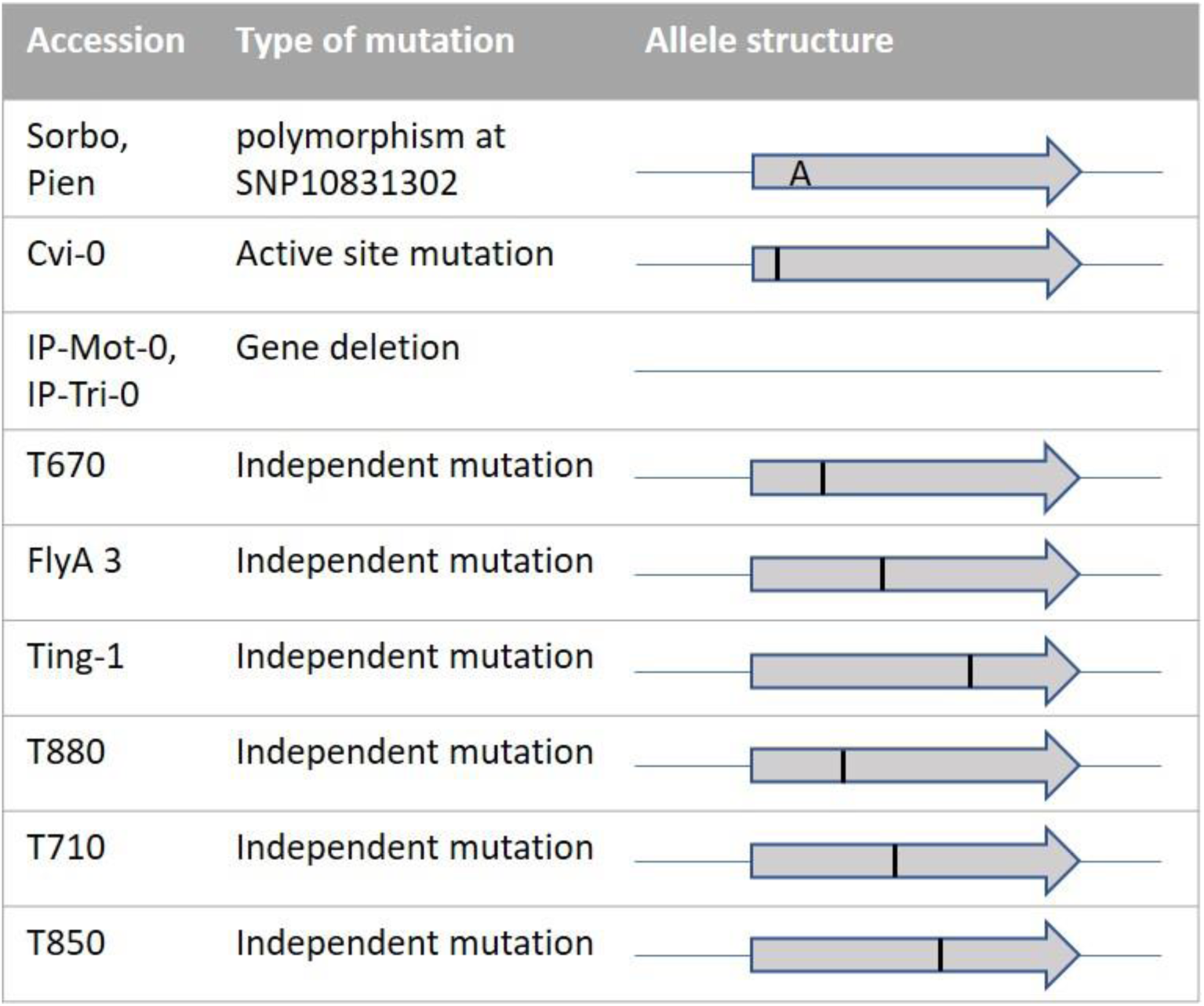
GS-OH structure. The structures of GS-OH in the 3-Butenyl accessions are illustrated. These mutations create premature stop codons.

| Accession | Type of mutation | Allele structure |
| --- | --- | --- |
| Sorbo, Pien | polymorphism at SNP10831302 |  |
| Cvi-0 | Active site mutation |  |
| IP-Mot-0, IP-Tri-0 | Gene deletion |  |
| T670 | Independent mutation |  |
| FlyA 3 | Independent mutation |  |
| Ting-1 | Independent mutation |  |
| T880 | Independent mutation |  |
| T710 | Independent mutation |  |
| T850 | Independent mutation |  |

**Figure 2:**
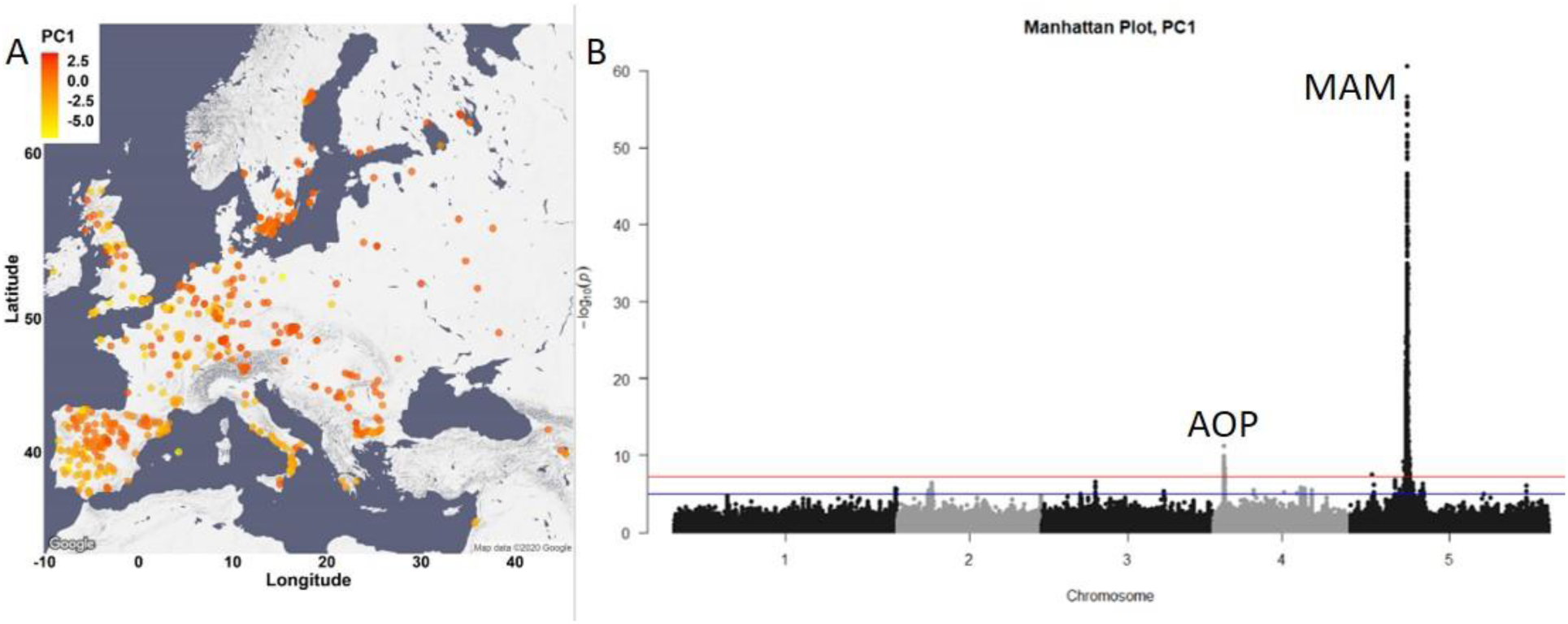
GSL variation across Europe is dominated by two loci. A. The accessions are plotted on the map based on their collection site, and colored based on their PC1 score. B. Manhattan plot of GWAS analyses using PC1. Horizontal lines represent 5% significance thresholds using Bonferroni (red) and permutations (blue).

To test which of the major causal loci are detectable in this collection and to identify new genomic regions that are associated with the observed GSL variation, we performed genome wide association (GWA, with EMMAX algorithms) analyses using the PC1 and PC2 values. This collection of natural accessions presents a dense variant map and is 3x larger than previous GSL GWA mapping populations. In spite of the large population size, both PC1 and PC2 based analyses identified the same two major peaks covering two of the known causal genes controlling GSL diversity (Figure 2B for PC1 GWA analyses, supp. Figure 2B for PC2 GWA analyses) (Brachi et al., 2015; Chan et al., 2011, 2010). The largest peak in both cases, is the GS-Elong locus on chromosome 5, containing the MAM1 (AT5G23010), MAM2 and MAM3 (AT5G23020) genes. The peak on chromosome 4 is the GS-AOP locus containing the AOP2 and AOP3 genes (AT4G03060 and AT4G03050, respectively). Previous F2, QTL and molecular experiments have shown that the genes within GS-AOP and GS-Elong loci are the causal genes for GSL variation within these regions (Benderoth et al., 2006; Brachi et al., 2015; Chan et al., 2011, 2010; D J Kliebenstein, Gershenzon, et al., 2001; D. Kliebenstein et al., 2002; Daniel J Kliebenstein et al., 2002; Kroymann & Mitchell-Olds, 2005; Pfalz et al., 2007; Wentzell et al., 2007). Surprisingly, none of the 8 other known natural variants within the GSL biosynthetic pathway were identified by GWA including three that were found with 96 accessions and three that were found with 595 accessions using PC1 and 2 (Brachi et al., 2015; Chan et al., 2011, 2010; Daniel J. Kliebenstein, 2009). It is possible that the extended sampling of accessions may have created genomic and demographic issues that influenced this high false-negative error rate where ∼80% of validated natural variants found using multiple RIL populations were missed.

### Complex GSL Chemotypic Variation

One potential complicating factor is that GSL chemotypic variation is best described as a discrete multimodal distribution involving the epistatic interaction of multiple genes which PCA’s linear decomposition cannot accurately capture (Figure 1). To test if PCA was inaccurately describing GSL chemotypic variation, we directly called the specific GSL chemotypes in each accession. Using Arabidopsis QTL mapping populations and GWA, we have shown that the GS-AOP, Elong and OH loci determine seven discrete chemotypes, 3MSO, 4MSO, 3OHP, 4OHB, Allyl, 3-Butenyl, 2-OH-3-Butenyl, that can be readily assigned from GSLs phenotypic data (Brachi et al., 2015; Chan et al., 2011, 2010; D J Kliebenstein, Gershenzon, et al., 2001). Using accessions with previously known chemotypes and genotypes, we developed a phenotypic classification scheme to assign the chemotype for each accession (Figure 3, for details see methods and supp. Figures 3-5, for structures see Figure 1 and supp. Table 1). Since the Aliphatic GSLs composition in the seeds reliably indicate the GSL structural composition in the other plant’s life stages and tissues, assigning a chemotype for each accession based on the seeds composition is expected to be highly stable across tissues of the same accession (Brown et al., 2003; Chan et al., 2011, 2010; D J Kliebenstein, Gershenzon, et al., 2001; D J Kliebenstein, Kroymann, et al., 2001). Most accessions were classified as 2-OH-3-Butenyl (27%) or Allyl (47%) with lower frequencies for the other chemotypes. Mapping the chemotypes on Europe showed that the PCA decomposition was missing substantial information on GSL chemotype variation (Figure 3). Instead of a continuous distribution across Europe, the chemotype classifications revealed specific geographic patterns. Central and parts of northern Europe were characterized by a high variability involving the co-occurrence of individuals from all chemotypes. In contrast, southern Europe, including the Iberian Peninsula, Italy and the Balkan, has two predominant chemotypes, Allyl or 2-OH-3-Butenyl, that are separated by a sharp geographic partitioning (Figure 3, and supp. Figure 6). The few accessions in southern Europe belonging to other chemotypes were all accessions previously identified as having genomes identical to accessions in central Europe, suggesting that they are likely stock center seed contaminations (The 1001 Genomes Consortium, 2016). Uniquely, Swedish accessions displayed a striking presence of almost solely Allyl chemotypes that was not mirrored on the eastern coast of the Baltic Sea (Finnish, Lithuanian, Latvian or Estonian accessions). Directly assigning GSL variation by discrete chemotypes provided a more detailed image not revealed by PCA decomposition. Further, the different chemotypic to geographic patterns suggests that there may be different pressures shaping GSL variation particularly when comparing central and southern Europe.

**Figure 3:**
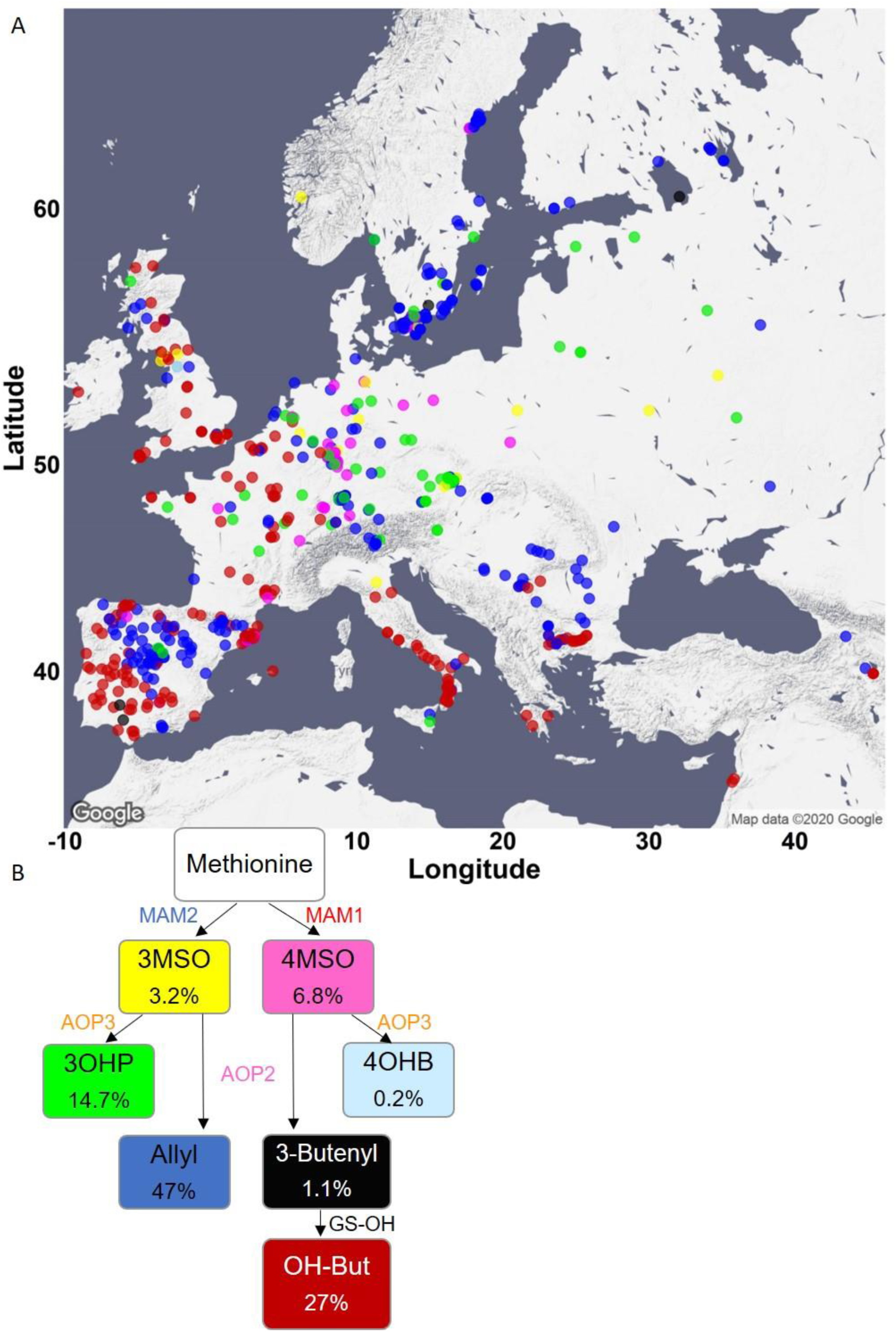
Phenotypic classification based on GSL content. A. Using the GSL accumulation, each accession was classified to one of seven aliphatic short chained GSL chemotypes based on the enzyme functions as follows: MAM2, AOP null: classified as 3MSO dominant, colored in yellow. MAM1, AOP null: classified as 4MSO dominant, colored in pink. MAM2, AOP3: classified as 3OHP dominant, colored in green. MAM1, AOP3: classified as 4OHB dominant, colored in light blue. MAM2, AOP2: classified as Allyl dominant, colored in blue. MAM1, AOP2, GS-OH non-functional: classified as 3-Butenyl dominant, colored in black. MAM1, AOP2, GS-OH functional: classified as 2-OH-3-Butenyl dominant, colored in red. The accessions were plotted on a map based on their collection sites and colored based on their dominant chemotype. B. The coloring scheme with functional GSL enzymes in the aliphatic GSL pathway is shown with the percentage of accessions in each chemotypes (out of the total 797 accessions) shown in each box.

### Geography and environmental parameters affect GSL variation

Because GSL chemotypes may be more reflective of local environment, we proceed to test if they are associated with weather parameters and landscape conditions. Further, given the difference in chemotype occurrence in central and southern Europe we hypothesized that these environmental connections may change between central and southern Europe. For these tests, we chose environmental parameters that capture a majority of the environmental variance and by that may describe the type of ecosystem (Ferrero-Serrano & Assmann, 2019). We assigned each accession the environmental value based on its location. These environmental parameters include geographic proximity (distance to the coast), precipitation descriptors (precipitation of wettest and driest month) and temperature descriptors (maximal temperature of warmest month and minimal temperature of coldest month) capture major abiotic pressures as well as provide information about the type of ecosystem in which each accession exists. We ran a multivariate analysis of variance (MANOVA) for each geographic area separately (north and central vs south, as shown in supp. Figure 6). This showed significant difference in how the GSL chemotypes associated to the environmental parameters across Europe. This was best illustrated by the two dominant chemotypes, Allyl and 2-OH-3-Butenyl, showing opposing relationships to the precipitation in the driest month. In Northern and Central Europe, the Allyl chemotype is more associated with lower precipitation in the driest month, while accessions with 2-OH-3-Butenyl as the dominant chemotype are associated with higher precipitation in the driest month. In Southern European accessions, this association is inverted (Figure 4A,B). This suggests that the relationship of GSL chemotype to environmental parameters vary across geographic regions of Europe rather than fitting a simple linear model.

**Figure 4:**
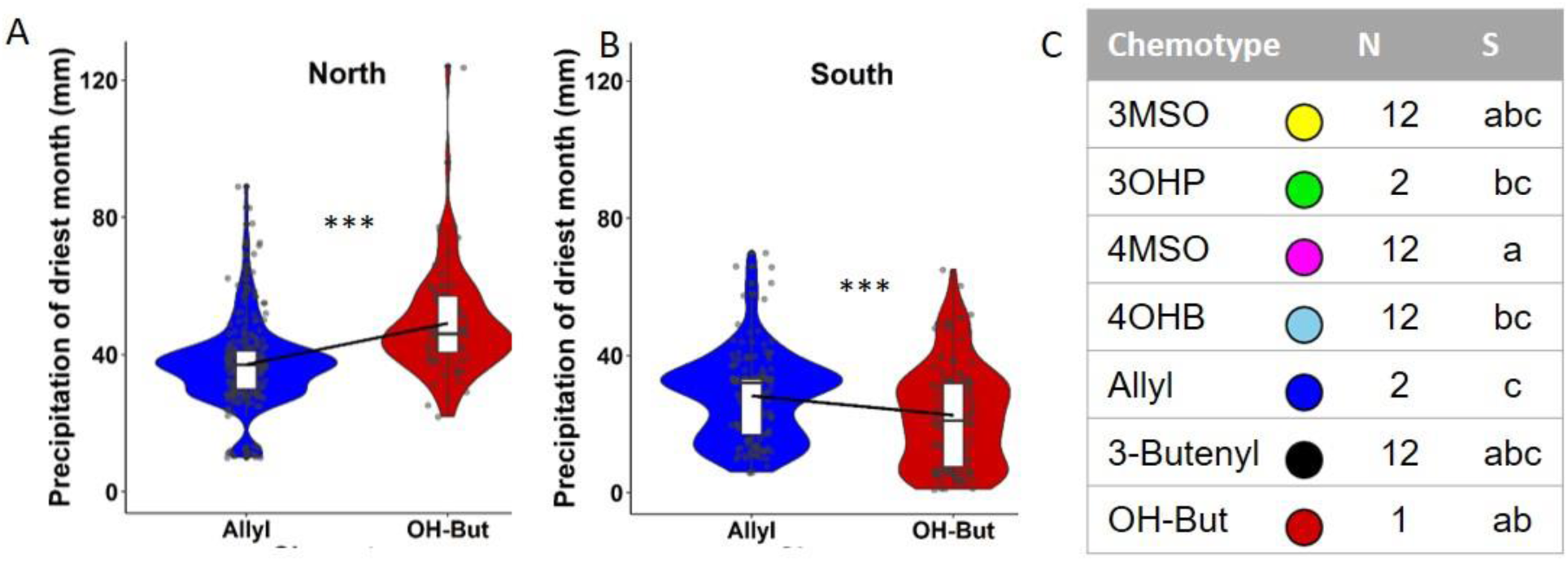
Environmental conditions differentially associate with GSLs across geographic location. A. B. The association of the two major chemotypes allyl and 2-OH-3-Butenyl to precipitation values of the driest month. Significance was tested by t-Test, P = 0.00000258 for the North (Slope= 0.01), P = 0.0005521 for the South (Slope= −0.007). C. MANOVA was performed for the south and north as indicated in methods section, followed by pairwise comparisons of least-squares means. Numbers indicate chemotypes with significant differences in the North and letters indicate chemotypes with significant differences in the south. OH-But= 2-OH-3-Butenyl.

As the two main chemotypes in the collection differ by the length of the carbon chain (C3 for Allyl, C4 for 2-OH-3-Butenyl), we created a linear model to further check the interaction between each environmental condition to geography in respect to the carbon chain length. Most of the environmental parameters significantly interacted with geography, meaning that the relationship of environment to GSL alleles change across geographic areas (supp. Figure 7, for details on the models see methods). Conducting this analysis for each of the geographic areas separately highlighted this by showing that these parameters have different effects on the carbon chain length in each of the areas (supp. Figure 7). This was true when the model was run with or without ancestral population state being included in the model (The 1001 Genomes Consortium, 2016).

Using a random forest machine learning approach provided similar results with different environment to chemotype relationships in the north and south (sup. Figure 8), supporting the hypothesis that the GSL chemotype to environment relationships change across regions within Europe.

### The genetic architecture of GSL variation

The presence of different GSL chemotype to environmental relationships across Europe raises the question of how these chemotypes are generated. Are these chemotypes from locally derived alleles or obtained by the intermixing of widely distributed causal alleles. Further, if there are multiple alleles, do they display within species convergent or parallel signatures. We focus on the GS-AOP, GS-Elong, and GS-OH loci, the causal genes creating Arabidopsis GSL chemotypes, and use the available genomic sequences in all of these accessions to investigate the allelic variation in these genes to map the allelic distribution and test the potential for convergent and/or parallel evolution within each locus.

GS-Elong: Because the variation in the GS-Elong locus is caused by complex structural variation in MAM1 and MAM2 that is not resolvable using the available data from short-read genomic sequence, we used the MAM3 sequence within this locus to ascertain the genomic relationship of accessions at the causal GS-Elong locus (Kroymann et al., 2003). We aligned the MAM3 sequence from each of the accessions, rooted the tree with the *Arabidopsis lyrata* orthologue (MAMc), and colored the tree tips based on the accessions dominant chemotype.

The accessions were distributed across eight distinctive clades with each clade clustering accessions having either a C3 or C4 status (Figure 5A). The clades C3/C4 status altered across the tree with three of the clades C3 dominant (MAM2 expressed), and five clades being C4 dominant (MAM1 expressed). Further supporting the use of MAM3 is that the accession assignments to these clades agree with available bacterial artificial chromosome-based sequencing of the GS-Elong region from 15 accessions (Figure 5B). While there are multiple functional alleles for both C3 and C4 chemotypes, the genomic sequence and phylogeny does not appear consistent with a simple parallel evolution model where one allele/population is the basis for the independent derivation of all alternative alleles. This is illustrated by the difference in the genomic arrangement of Clade 4 and 5 which both create C4 GSLs. Clade 4 has a copy of MAM2 and MAM1 while Clade 5 has two copies of MAM1 (Figure 5). It appears that Clade 4 is the basis for two independent C3 alleles via separate deletions of MAM1 (Clades 1 and 3) and a separate C4 allele via a deletion of MAM2 (Clade 2, Figure 5). Unfortunately, no long-read sequencing is available in accessions from Clade 6 or 7 and locus-specific de novo alignment of short-read sequences in these accessions was not able to resolve the regions complexity. Filling in these clades would be necessary to better understand convergent/parallel events giving rise to GSL chemotypes.

**Figure 5:**
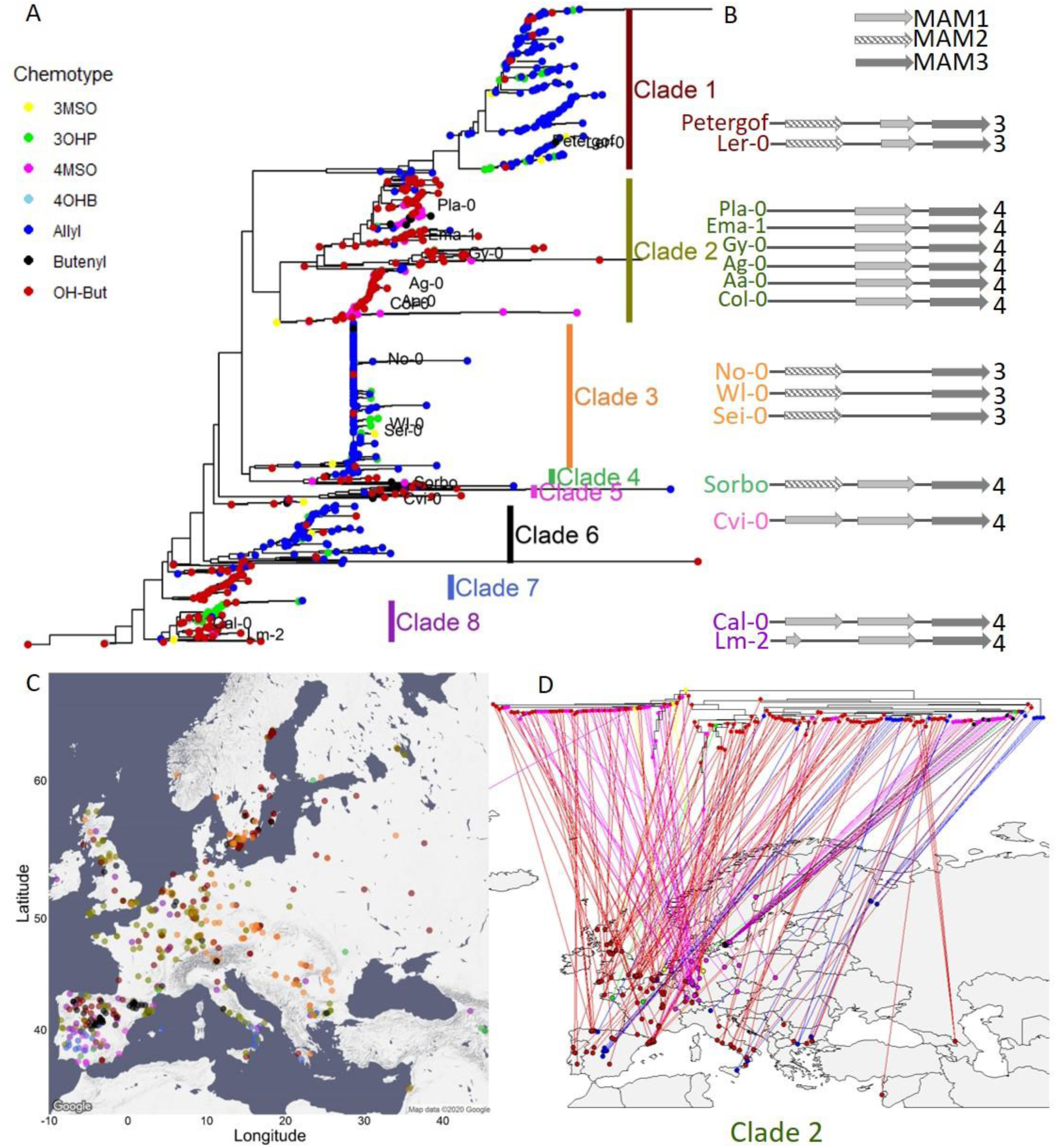
MAM3 phylogeny. A. MAM3 phylogeny of *Arabdopsisi thaliana* accessions, rooted by *Arabidopsis lyrata* MAMc, that is not shown because of distance. Tree tips are colored based on the accession chemotype. The named accessions indicate that GSL-Elong region of these accessions was previously sequenced (Kroymann et. al. 2003). B. The genomic structure of the GSL-Elong regions in the previously sequenced accessions is shown based on Kroymann et. al. 2003. The accession names are colored based on their clades. The color of the name of the accession indicates the clade it belongs. Bright grey arrows represents MAM1 sequences, dashed arrows represents MAM2 sequences. Dark grey arrows represent MAM3 sequences. The number to the right of the genomic cartoon represents the number of carbons in the side chain. C. Collection sites of the accessions, colored by their clade classification (from section A). D. Clade 2 reflection on the map.

**Figure 6:**
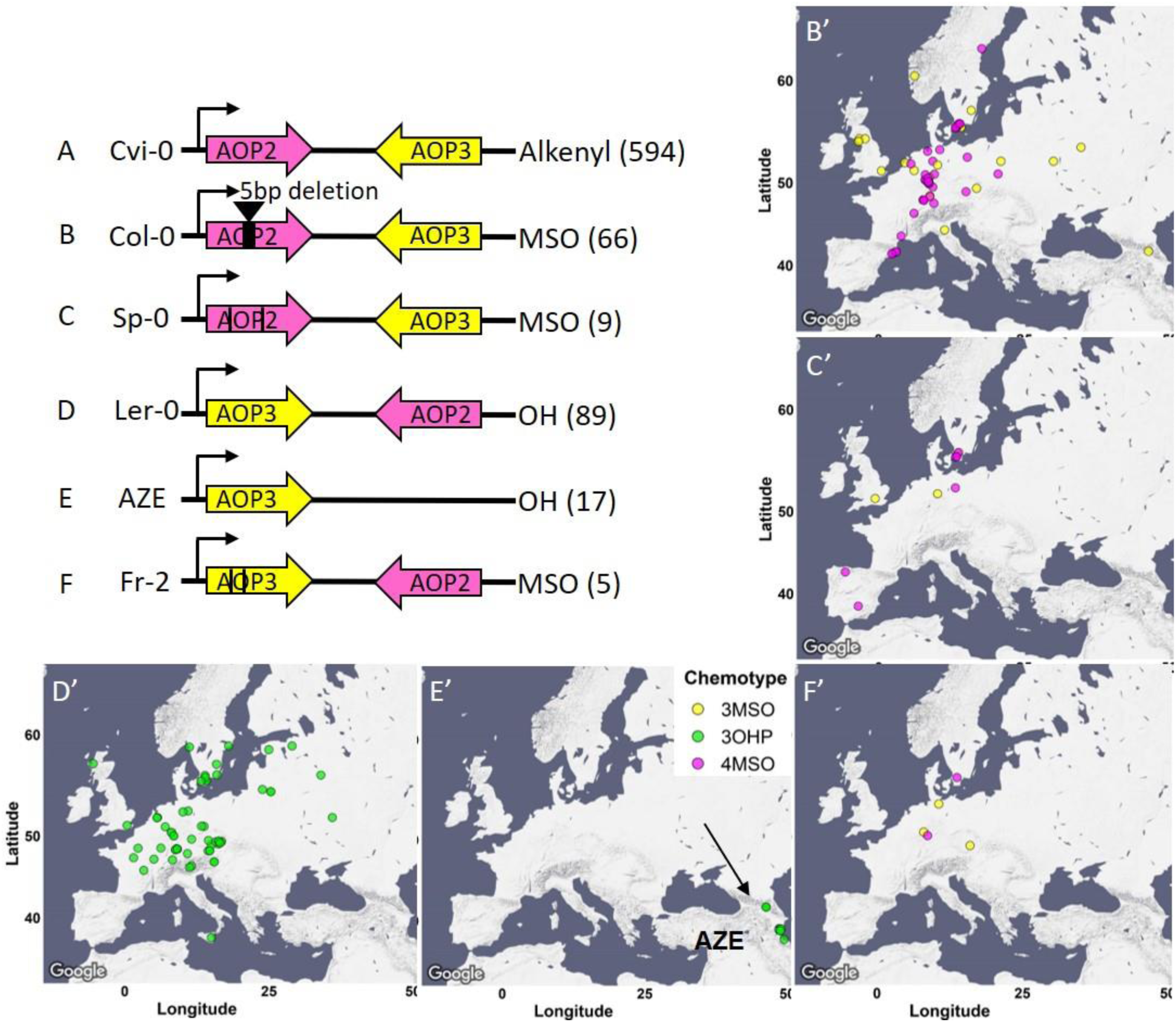
AOP Genomic structure. The genomic structure and causality of the major AOP2/AOP3 haplotypes are illustrated. Pink arrows show the AOP2 gene while yellow arrows represent AOP3. The black arrows represent the direction of transcription from the AOP2 promoter as defined in the Col-0 reference genome. Its position does not change in any of the regions. The black lines in Sp-0 and Fr-2 presence the position of independent variants creating premature stop codons. The GSL chemotype for each haplotype is listed to the right with the number of the accessions in brackets. The maps show the geographic distribution of the accessions from each structure.

Interestingly, the most basal clade has no copy of MAM2, raising the question of where MAM2 arose (Figure 5). This suggests that true ancestral state(s) of this locus is not represented in this collection and would need to be searched for in other populations of *Arabidopsis thaliana*, if it exists in extant populations.

Using this phylogeny, we investigated the presence of the different GS-Elong haplotypes across Europe to ask if each region has a specific allele/clade or if the alleles are distributed across the continent. Specifically, we were interested if the strong C3/C4 partitioning in southern Europe was driven by the creation of local alleles or if this partitioning might contain a wide range of alleles. If the latter is true, this would argue for a selective pressure shaping this C3/C4 divide. We plotted the accessions on the map and colored them based on their GS-Elong clade (Figure 5C). This showed that the strong C3/C4 partition in the Iberian Peninsula contains haplotypes from all the GS-Elong clades except Clade 3 and is not caused by local alleles. This suggests that the strong geographic partitioning of the C3/C4 chemotypes in Iberia may be driven by selective pressure causing the partitioning of the chemotypes rather than neutral demographic processes.

Shifting focus to all of Europe showed that while most clades were widely distributed across Europe there were a couple over-arching patterns (Figure 5C, and supp. Figure 9). GS-Elong clades 1 and 6 follow a pattern that fits with alleles located within the Iberian glacial refugia that then moved north. In contrast the absence of clade 3 from Iberia is more parsimonious with a glacial refugia in the Balkans followed by a northward movement wherein it mixed with the other clades. Other clades never moved north and are exclusive to the south as shown by clades 5 and 7. While these are both C4 clades, other C4 clades like clades 2 and 8 were able to move north (Figure 5D, and supp. Figure 9, respectively). This suggests that there are either differences in their GSL chemotype influencing their distribution or there are neighboring genes known to be under selection in Arabidopsis like FLC (AT5G10140) that may have influenced their distribution. In combination, this suggests that a complex demography is involved in shaping the chemotypes identity with some regions, Iberia, showing evidence of local selection while other regions, central Europe, possibly showing a blend requiring further work to delineate (supp. Figure 9).

GS-AOP: Side chain modification of the core MSO GSL is determined by the GS-AOP locus. Most of the accessions contain a copy of AOP2 and a copy of AOP3, but only one of them will be functionally expressed (Chan et al., 2010), while in some cases both will be nonfunctional. To better understand the demography and evolution of the GS-AOP locus, we separately aligned the AOP2 and AOP3 sequences, rooted each tree with the *Arabidopsis lyrata* orthologue, and colored the trees tips based on the accessions dominant chemotype.

The phylogenetic trees shared a very similar topology, yielding a clear separation between alkenyl (AOP2 expressed) and hydroxyalkyl (AOP3 expressed) accessions. Alkenyl expressing accessions like Cvi-0 with an expressed copy of the AOP2 enzyme formed a single contiguous cluster (Figure 6A). In contrast, hydroxyalkyl accessions clustered into two separate groups with one group of 3OHP dominant accessions partitioning from the rest of the accessions at the most basal split in the tree (supp. Figure 10). This haplotype is marked by having an inversion swapping the AOP2 and AOP3 promoters as shown in bacterial artificial chromosome sequencing of the Ler-0 accession (Figure 6D) (Chan et al., 2010). The tree also identified a second group of 3OHP dominant accessions located among the alkenyl accessions. Analyzing the sequences of these accessions reveals that this small group of 3OHP accessions have a complete deletion of AOP2 and contain only AOP3 (Figure 6E). Thus, there are at least two independent transitions from Alkenyl to Hydroxyalkyl GSLs within Arabidopsis, neither of which are related to the Alkenyl to Hydroxyalkyl conversion within *Arabidopsis lyrata*.

The null accessions (MSO dominant chemotypes) were identifiable in all the major clades on the tree (supp. Figure 10, middle column of heatmap) suggesting that there are independent LOF mutations that abolish either AOP2 or AOP3. Deeper examination of the sequences of these accessions identified three convergent LOF alleles leading to the MSO chemotype. Most of the null accessions harbor a 5 bps deletion in their AOP2 sequence, that causes a frameshift mutation. This mutation arose within the Alkenyl haplotype and was first reported in the Col-0 reference genome (Figure 6B) (D J Kliebenstein, Lambrix, et al., 2001). In addition, there are additional independent LOF events arising in both the alkenyl haplotype (e.g. Sp-0, Figure 6C), and within the Ler-0 inversion haplotype (e.g. Fr-2, Figure 6F). Thus, GS-AOP has repeated LOF alleles arising within all the major AOP haplotypes suggesting convergent evolution of the MSO chemotype out of both the Alkenyl and Hydroxyalkyl chemotypes.

Using the combined chemotype/genotype assignments at GS-AOP, we investigated the distribution of the alleles across Europe. The Alkenyl haplotype is spread across the entire continent. In contrast, the hydroxyalkyl haplotypes are more local. The Ler-like 3OHP haplotype is present in only central and north Europe (Figure 6D), while the other 3OHP haplotype, possessing only AOP3, is limited to Azerbaijan, along the Caspian Sea (Figure 6E). In contrast to the distinct hydroxyalkyl locations, the distribution of the independent LOF null haplotypes overlaps with all of them being located within central and north Europe (Figure 6B, C and F). The fact that these independently derived LOF alleles are all contiguous suggests that there may be a benefit to these alleles specific to Central Europe.

GS-OH: The final major determinant of natural variation in Arabidopsis GSL chemotype is the GS-OH enzyme that adds a hydroxyl group to the 2 carbon on 3-butentyl GSL to create 2-OH-3-butenyl GSL. Previous work had suggested two GS-OH alleles measurable in the seed, a functional allele in almost all accessions and a non-functional allele caused by active site mutations represented by the Cvi-0 accession (Hansen et al., 2008). Because of functional epistasis, we can only obtain functional phenotypic information from accessions that accumulate the GS-OH substrate, 3-butenyl GLS. This identified 11 accessions with a non-functional GS-OH. Surveying these 11 accessions in the polymorph database identified multiple independent LOF events. One of these 11 accessions have the Cvi active site mutations, two accessions have a shared nonsense SNP that introduce premature stop codons, and two accessions have a complete loss of this gene (Table 1). We could not identify the causal LOF allele in the other six accessions due to sequence quality in the databases. All of these independent GS-OH LOF alleles are phylogenetically positioned within groups of accessions that largely do not accumulate 3-butenyl GLS, e.g. 3 carbon or non-alkenyl accessions suggesting that the functional epistasis may be influencing the generation of these alleles. Thus, we searched the accessions that do not accumulate 3-butenyl GLS and have effectively hidden the GS-OH function for these GS-OH LOF events (Supp table 2). In each case, the LOF allele is more frequent in the non 4 carbon-alkenyl accessions than expected by random chance. This suggests that there is a bias against 3-butenyl GSL synthesis as the LOF alleles are more frequent when the GS-OH gene is hidden by functional epistasis. This agrees with the fact that the 3-butenyl chemotype is the most sensitive to generalist lepidopteran herbivory (Hansen et al., 2008). Thus, these mutations may represent ongoing pseudogenization of the GS-OH gene when it is functionally hidden by epistasis at the GS-AOP and GS-Elong loci. These LOF events would then only be displayed upon rare admixture with 2-OH-3-Butenyl accessions.

## Discussion

Understanding the genetic, demographic and environmental factors that shape variation within a trait in a population is key to understanding trait evolution. In this work we used Aliphatic GSLs in seeds of Arabidopsis thaliana to query how genetics, geography, environment and demography intersect to shape chemotypic variation across Europe. We found that environmental conditions, together with geography affect the presence and distribution of chemotypes within the accessions. This was demonstrated by specific traits that were associated with specific environmental conditions, and this association was shifted across the continent. Comparing the associations of traits to specific environmental conditions in central Europe versus the south revealed different, sometimes even inverse, behaviors. For example, In the Iberian Peninsula, 2-OH-3-Butenyl was positively associated with potential drought while in Central Europe, it was the opposite GS-Elong allele showing association. This showed that chemotypic variation across Europe is created by a blend of all these processes that differ at the individual loci and required the simultaneous analysis of genotype and phenotype to fully interpret.

In contrast to the bimodal distribution in the Aliphatic GSL traits, each of the three major Aliphatic GSL loci showed allelic heterogeneity with multiple independent structural variants that recreated the same phenotypic variation. The GS-AOP locus had numerous events with the AOP3 variant of GS-AOP arising via at least two independent events and the Null allele being generated at least 15 independent times. The GS-AOP null alleles convergently arose from all the different functional haplotypes. It is less clear if the independent GS-AOP AOP3 alleles should be classified as convergent or parallel due to a lack of clarity in what is the ancestral state. Similar to the independent GS-AOP null alleles, there were numerous independent GS-OH LOF variants with at least 9 independent events. While these are parallel GS-OH LOF events because they came from a single functional GS-OH group, their ability to accumulate depends on the epistatic silencing of GS-OH by the GS-AOP and GS-Elong loci. The GS-Elong locus also had an extensive level of allelic heterogeneity hallmarked by a shifting expression of the MAM1 or MAM2 gene, again with hallmarks of both parallel and convergent processes. Interestingly, at both the GS-AOP and GS-Elong loci, one gain-of-function event (e.g. AOP3 in GS-AOP locus, and MAM2 in GS-Elong locus) is concurrently linked to a loss-of-function of the other gene at the locus (AOP2 and MAM1, respectively). These structural variants are shaped such that the chemotypes show distinct separations without any intermediate phenotypes. This allelic heterogeneity is in contrast to previous work on other biotic interactions genes like pathogen resistance gene-for-gene loci that typically have two moderate frequency stable alleles creating the phenotypic variation within the species (Atwell et al., 2010; Corrion & Day, 2001; MacQueen, Sun, & Bergelson, 2016). In other cases alleles of genes involved in biotic defense can present more complex patterns, e.g. natural variation in the immune gene *ACCELERATED CELL DEATH 6* (ACD6) is caused by a rare allele causing an extreme lesion phenotype. It is not yet clear what selective pressures influence ACD6 genetic variation (Todesco et al., 2010; Zhu et al., 2018). The contrast where Aliphatic GSL loci have high levels of allelic heterogeneity for independent and recurrent LOF and GOF events while other resistance genes have more stable biallelic variation suggests that there are different selective regimes influencing these loci. Further work is needed to assess the range of allelic heterogeneity in loci controlling resistance to diverse biotic traits within the environment.

The allelic heterogeneity at these loci illustrates the benefit of simultaneously tracking the phenotype and genotype when working to understand the distribution of trait variation. For example, the Iberian Peninsula and the Mediterranean had low variability in Aliphatic GSL chemotype with the chemotypes not overlapping while central/north Europe had high Aliphatic GSL diversity with the chemotypes overlapping. At first glance, this contrasts with previous work showing that the Iberian Peninsula and the Mediterranean are more genetically diverse. However, this discrepancy was caused by one of the causal loci. Specifically, the GS-AOP locus is largely fixed as the Alkenyl allele in Iberia/Mediterranean with the alternative GS-AOP alleles enriched in central Europe. In contrast to GS-AOP, Iberia and the Mediterranean were highly genetically diverse for the GS-Elong locus and appear to contain all the variation in GS-Elong found throughout Europe (The 1001 Genomes Consortium, 2016). Thus, the chemotypic divergence from genomic variation expectations was driven by just the GS-AOP locus. This indicates that the high level of chemotypic variation in central Europe is a blend of alleles that moved from the south (GS-Elong) and alleles that possibly arose locally (GS-AOP, both nulls and AOP3). Further, the chemotypes found in any one region appear to be created by a combination of alleles moving across the continent, local generation of new polymorphisms and local selective pressures that shape the chemotypes distribution across the landscape.

One difficulty in interpreting the evolutionary processes, e.g. parallel v convergent, especially for structural variants illustrated by all the three loci is the complication in properly identifying the ancestral state of the population. While this could typically be done by relying on shared loci with sister species, this is not possible in this case as *Arabidopsis lyrata* and *halleri* have genetic variation at GS-Elong and GS-AOP creating the exact same phenotypes. Further, neither of these sister species have yet been found to have a functional GS-OH (Heidel, Clauss, Kroymann, Savolainen, & Mitchell-Olds, 2006; Ramos-Onsins, Stranger, Mitchell-Olds, & Aguadé, 2004; Windsor et al., 2005). The MSO chemotypes could be viewed as convergent evolution within a species, as the MSO phenotype independently re-occurred multiple times in the AOP2 and AOP3 genetic backgrounds. However, it is not clear how to classify the different AOP3 (AZE v Ler) types as it not clear if the AOP2 or AOP3/Ler haplotype is ancestral within *Arabidopsis thaliana*. Another option to calling ancestral state is deep sampling in the species but even with these accessions, we do not appear to have reached the necessary threshold. For example previous work at the GS-Elong locus had suggested that the Sorbo accession, collected from Tajikistan, was the most likely ancestral state as it had a copy of both MAM1 and MAM2 (Kroymann et al., 2003). However, the phylogeny with this larger collection of accessions suggested that Sorbo is not ancestral. Further, a recent phylogeny of MAM genes across the Brassicales suggests that MAM2 is an *Arabidopsis thaliana* specific gene with an undefined origin (Abrahams et al., 2020). This suggests that to get a better understanding of the ancestral state to define evolutionary processes, especially for loci with allelic heterogeneity and structural variants, we need to broaden our phylogenetic context by deeper sampling within and between species.

Another complication caused by the allelic heterogeneity and differential selective pressures displayed within this system is that we were unable to detect a number of known and validated natural variants that are causal within this population. Specifically, the GWAS with this collection of 797 accessions was unable to find 80% of the known causal loci including one of the three major effect loci, GS-OH. Maximizing the number of genotypes and the SNP marker density was unable to overcome the complications imposed by the complex pressures shaping the distribution of these traits. In this system, the optimal path to identifying the causal polymorphisms has instead been a small number of Recombinant Inbred Line populations derived from randomly chosen parents. In complex adaptive systems, the optimal solution to identifying causal variants is likely a blend of structured mapping populations and then translating the causal genes from this system to the GWAS results and tracking the causal loci directly.

In this work we combined different approaches to uncover some of the parameters shaping the Aliphatic GSL content across Europe. Widening the size of the population will enable us to deepen our understanding on the evolutionary mechanisms shaping a phenotype in a population.

## Methods

### Plant materiel

Seeds for 1135 Arabidopsis (*Arabidopsis thaliana*) genotypes were obtained from the 1001 genomes catalog of Arabidopsis thaliana genetic variation (https://1001genomes.org/). All Arabidopsis genotypes were grown at 22°C/24°C (day/night) under long-day conditions (16-h of light/8-h of dark). Two independent experiments were performed, each of them included the full set of genotypes. In the analyses only accessions from Europe and around Europe were included (Figure 2A), resulting in an analysis of 797 accessions. List of the accessions can be found in supp. Table 1.

### GSL extractions and analyses

GSLs were measured as previously described (D J Kliebenstein, Gershenzon, et al., 2001; D J Kliebenstein, Kroymann, et al., 2001; D J Kliebenstein, Lambrix, et al., 2001). Briefly, ∼3mg of seeds were harvested in 200 μL of 90% methanol. Samples were homogenized for 3 min in a paint shaker, centrifuged, and the supernatants were transferred to a 96-well filter plate with DEAE sephadex. The filter plate with DEAE sephadex was washed with water, 90% methanol, and water again. The sephadex bound GSLs were eluted after an overnight incubation with 110μL of sulfatase. Individual desulfo-GSLs within each sample were separated and detected by HPLC-DAD, identified, quantified by comparison to standard curves from purified compounds, and further normalized to the weight. List of GSLs and their structure are in supplementary table Row GSLs data are in supplementary table 1B.

### Statistics, heritability, and data visualization

Statistical analyses were conducted using R software (https://www.R-project.org/) with the RStudio interface (http://www.rstudio.com/). For each independent GLS, a linear model followed by ANOVA was utilized to analyze the effect of accession, replicate, and location in the experiment plate upon the measured GLS amount. Broad-sense heritability (supplementary table 1C) for the different metabolites was estimated from this model by taking the variance due to accession and dividing it by the total variance. Estimated marginal means (emmeans) for each accession were calculated for each metabolite from the same model using the package emmeans (“CRAN - Package emmeans,” n.d.) (supplementary table 1D). Principal component analyses were done with FactoMineR and factoextra packages (Abdi & Williams, 2010). Data analyses and visualization was done using R software with tidyverse (Wickham et al., 2019) and ggplot2 (Kahle & Wickham, 2013) packages.

Principal component analyses were done with FactoMineR and factoextra packages (Abdi & Williams, 2010).

Maps were generated using ggmap package (“https://journal.r-project.org/archive/2013-1/kahle-wickham.pdf,” n.d.).

### Phenotypic classification based on GSL content

For each accession the expressed enzyme in each of the following families was determined based on the content (presence and amounts) of short chained Aliphatic GSLs:

MAM enzymes: the total amount of 3 carbons GSLs and 4 carbons GSLs was calculated for each accession. 3 carbons GSLs include 3MT, 3MSO, 3OHP and Allyl GSL. 4 carbons GSLs include 4MT, 4MSO, 4OHB, 3-butenyl and 2-OH-3-butenyl GSL (for structures and details see supp. Table 1). Accessions that the majority of Aliphatic short chained GSL contained 3 carbons in their side chains classified as MAM2 expressed (supp. Figure 3). Accessions that the majority of Aliphatic short chained GSL contained 4 carbons in their side chains classified as MAM1 expressed (supp. Figure 3). The accessions were plotted on a map based on their original collection sites (supp. Figure 3).

AOP enzymes: the relative amount of alkenyl GSL, alkyl GSL and MSO GSL were calculated in respect to the total short chained Aliphatic GSL as follows:

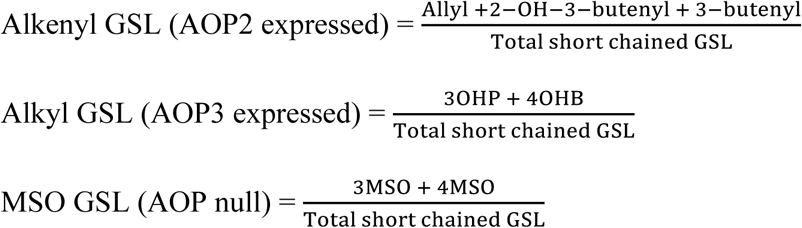

The expressed AOP enzyme was determined based on those ratios: accessions with majority alkenyl GSL were classified as AOP2 expressed. Accessions with majority of alkyl GSL were classified as AOP3 expressed. Accessions with majority of MSO GSL were classified as AOP null. The accessions were plotted on a map based on their original collection sites (supp. Figure 4).

GS-OH enzyme: the ratio between 2-OH-3-butenyl GSL to 3-butenyl GSL was calculated only for MAM1 expressed accessions (accessions that the majority of GSLs contain 4 carbons in their side chain). Accessions with high amounts of 2-OH-3-butenyl GSL were classified as GS-OH functional. Accessions with high amounts of 3-butenyl GSL were classified as GS-OH non-functional. The accessions were plotted on a map based on their original collection sites (supp. Figure 5).

Each accession was classified to one of seven Aliphatic short chained GSLs based on the combination of the dominancy of the enzymes as follows: MAM2, AOP null: classified as 3MSO dominant. MAM1, AOP null: classified as 4MSO dominant. MAM2, AOP3: classified as 3OHP dominant. MAM1, AOP3: classified as 4OHB dominant. MAM2, AOP2: classified as Allyl dominant. MAM1, AOP2, GS-OH non-functional: classified as 3-Butenyl dominant. MAM1, AOP2, GS-OH functional: classified as 2-OH-3-Butenyl dominant. The accessions were plotted on a map based on their original collection sites and colored based on their dominant chemotype (Figure 3).

### Environmental data

Environmental data was obtained from the 1001 genomes website (https://1001genomes.org/, for geographical data) and from the Arabidopsis CLIMtools (http://www.personal.psu.edu/sma3/CLIMtools.html, (Ferrero-Serrano & Assmann, 2019)) for environmental data. We used the five variables that captured a majority of the variance in this dataset including maximal temperature of warmest month (WC2_BIO5), minimal temperature of coldest month (WC2_BIO6), precipitation of wettest month (WC2_BIO13), precipitation of driest month (WC2_BIO14), and distance to the coast (in Km).

### Environmental MANOVA

Linear models to test the effect of geographical and environmental parameters (supp. Figure 1, 8) were conducted using dplyr package (“CRAN - Package dplyr,” n.d.) and included the following parameters:

Supp. Figure 1-linear models for collection sites: PC score ∼ Latitude + Longitude + Latitude * Longitude.

Supp. Figure 7 - for all the data: C length (C3 or C4) ∼ Genomic group + Geography (north versus south) +Max temperature of warmest month+ Min temperature of coldest month+ Precipitation of wettest month+ Precipitation of driest month+ Distance to the coast + Geography *Genomic group + Geography * Max temperature of warmest month + Geography * Min temperature of coldest month+ Geography * Precipitation of driest month+ Geography * Precipitation of wettest month + Geography *Distance to the coast.

For the north and the south: C length (C3 or C4) ∼ Genomic group + Geography (north versus south) +Max temperature of warmest month+ Min temperature of coldest month+ Precipitation of wettest month+ Precipitation of driest month+ Distance to the coast.

Multivariate analysis of variance (MANOVA) models to check the effect of environmental variables on the chemotype identity for each collection (Figure 4 and supp. Figure 7) included the following parameters:

Max temperature of warmest month+ Min temperature of coldest month+ Precipitation of wettest month+ Precipitation of driest month+ Distance to the coast∼ Chemotype.

### Random Forest analyses

Random forest analyses was conducted using the “randomForest” and “ElemStatLearn” packages in Rstudio (“CRAN - Package ElemStatLearn,” n.d., “CRAN: R News,” n.d.; Liaw & Wiener, 2002). In these analyzes we used the environmental parameters and genomic group data to predict the chemotype identity, after excluding the low frequencies chemotypes (4OHB and 3-Butenyl from all of them, 3MSO from the south).

### Genome wide association studies

The phenotypes for GWAS were each accession value for PC1 and 2. GWAS was implemented with the easyGWAS tool (Grimm et al., 2017) using the EMMAX algorithms (Kang et al., 2010)and a minor allele frequency (MAF) cutoff of 5%. The results were visualized as manhattan plots using the qqman package in R (Turner, 2014).

### Phylogeny

Genomic sequences from the accessions for MAM3 – AT5G23020, AOP2 – Chr4, 1351568 until 1354216, AOP3 - AT4G03050.2, and GS-OH – AT2G25450 were obtained using the Pseudogenomes tool (https://tools.1001genomes.org/pseudogenomes/#select_strains).

Multiple sequence alignment was done with the msa package in R, using the ClustalW, ClustalOmega, and Muscle algorithms (Bodenhofer, Bonatesta, Horejš-Kainrath, & Hochreiter, 2015). Phylogenetic trees were generated with the ggtree package in R (Yu, 2020). Each tree was rooted by the genes matching *Arabidopsis layrata’s* functional orthologue or closest homologue.

## Acknowledgments

This work was supported by the National Science Foundation, Directorate for Biological Sciences, Division of Molecular and Cellular Biosciences (grant no. MCB 1906486 to D.J.K) and Division of Integrative Organismal Systems (grant no. IOS 1655810 to D.J.K, and IOS 1754201 to R.A), and by the United States-Israel Binational Agricultural Research and Development Fund (to D.J.K. and E.K., grant no. FI–560–2017). We thank Dr. Allison Gaudinier (Department of Plant and Microbial Biology, University of California, Berkeley), Dr. Tobias Züst (Institute of Plant Sciences, University of Bern), Dr. Christopher W. Wheat (Department of Zoology, Stockholm University) and Dr. Daniel Runcie (Department of Plant Sciences, University of California Davis.) for critical reading of the manuscript.

