## Supplemental figures for "Genetic variation, environment and demography intersect to shape Arabidopsis defense metabolite variation across Europe"

**Supplementary figures**

Supplementary table 1: A. List of GSLs and structures. B. Accessions and GSL data – raw data. C. Heritability values. D. Accessions and GSL data – emmeans.

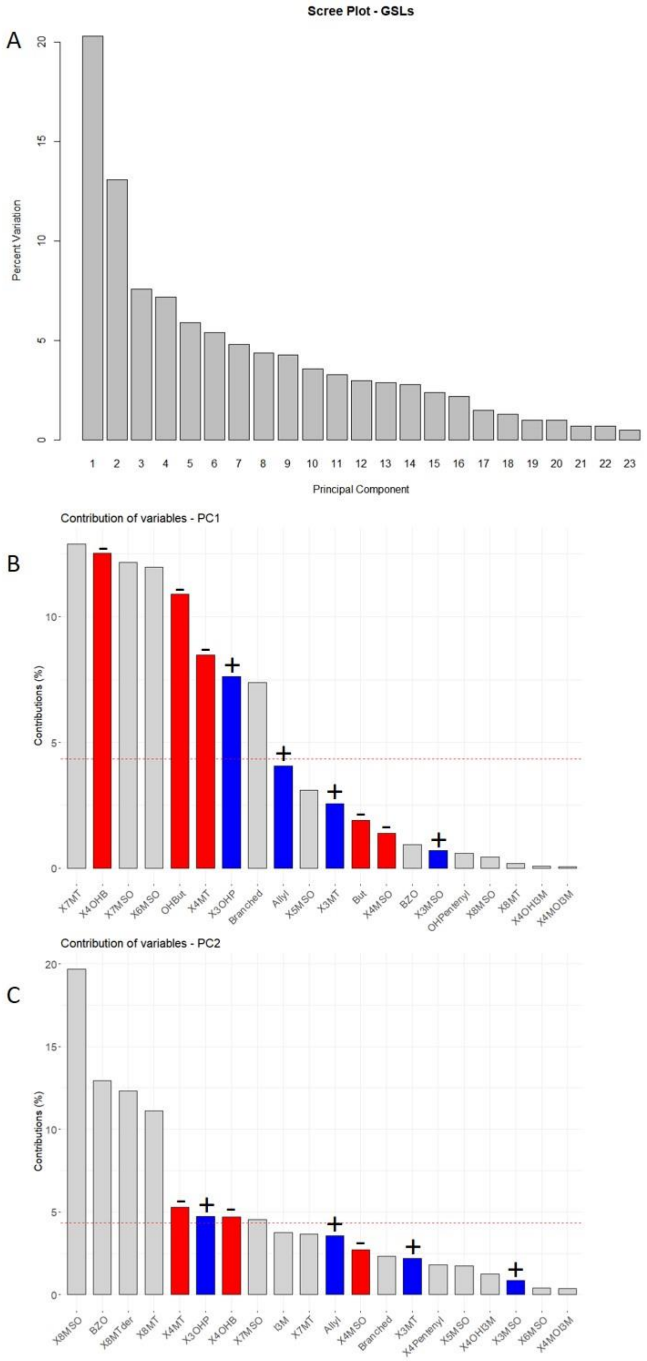


D

| Response: PC1 | | | | | | |
|---|---|---|---|---|---|---|
| | Df | SumSq | MeanSq | F | value | Pr(>F) |
| Lat | 1 | 67.3 | 67.31 | 16.368 | 5.73E-05 | *** |
| Long | 1 | 302.2 | 302.163 | 73.478 | 2.20E-16 | *** |
| Lat:Long | 1 | 92.4 | 92.45 | 22.481 | 2.52E-06 | *** |
| Residuals | 793 | 3261.1 | 4.112 | | | |
| **Response: PC2** | | | | | | |
| | Df | SumSq | MeanSq | F | value | Pr(>F) |
| Lat | 1 | 6.45 | 6.452 | 2.3147 | 0.1286 | |
| Long | 1 | 89.22 | 89.217 | 32.0069 | 2.15E-08 | *** |
| Lat:Long | 1 | 100.05 | 100.054 | 35.8949 | 3.16E-09 | *** |
| Residuals | 793 | 2210.43 | 2.787 | | | |

Supplementary figure 1: **GSL based PC analysis.** A. Percentage of variance explained by each principal component. B, C. contribution of the individual GSLs to PC1 (B) and PC2 (C). Red bars: contribution of 4 carbon GSLs, blue bars: contribution of 3 carbons GSLs. +/- above the bar indicates if the contribution of the variable is positive or negative. D. Linear model for PC1 and PC2 scores with the geographic parameters. Lat= Latitude, Long= Longitude.

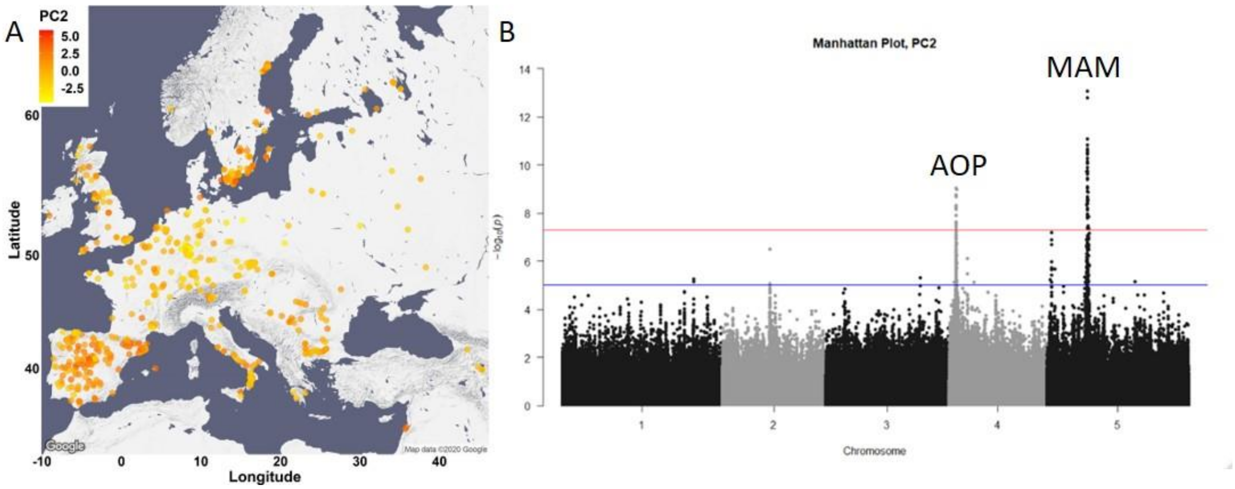


Supplementary figure 2: **GSL variation across Europe is domninated by two loci.** A. The accessions were plotted on the map based on their collection site, and colored based on their PC2 score. B. Manhattan plot of GWAS analyses using PC2. Horizontal lines represent 5% significance thresholds using Bonferroni (red) and permutations (blue).

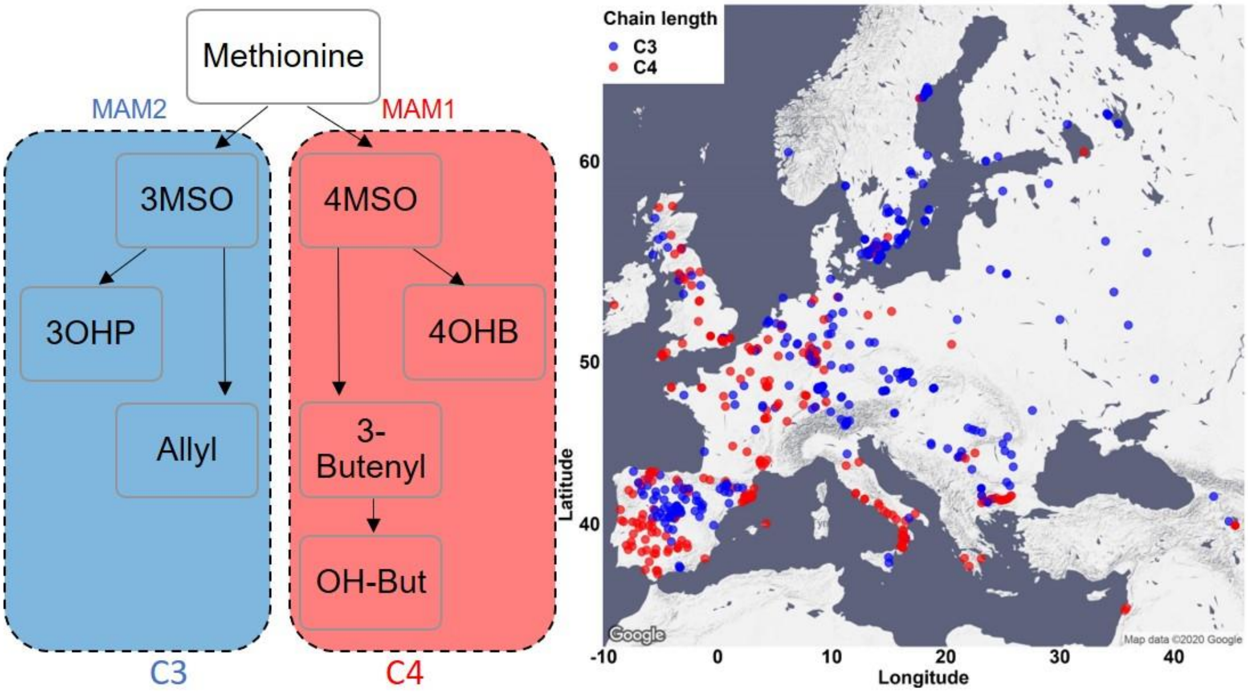


Supplementary figure 3: **Phenotypic classification based on the dominant MAM enzyme.** Accessions were classified based on the side chain length of the aliphatic short chained GSLs. Accessions with a majority of GSLs containing 3 carbons in their side chains are classified as MAM2 dominant, and colored in blue. Accessions with the majority of aliphatic short chained GSLs containing 4 carbons in their side chains are classified as MAM1 dominant, and colored in red. The accessions were plotted on a map based on their original collection sites.

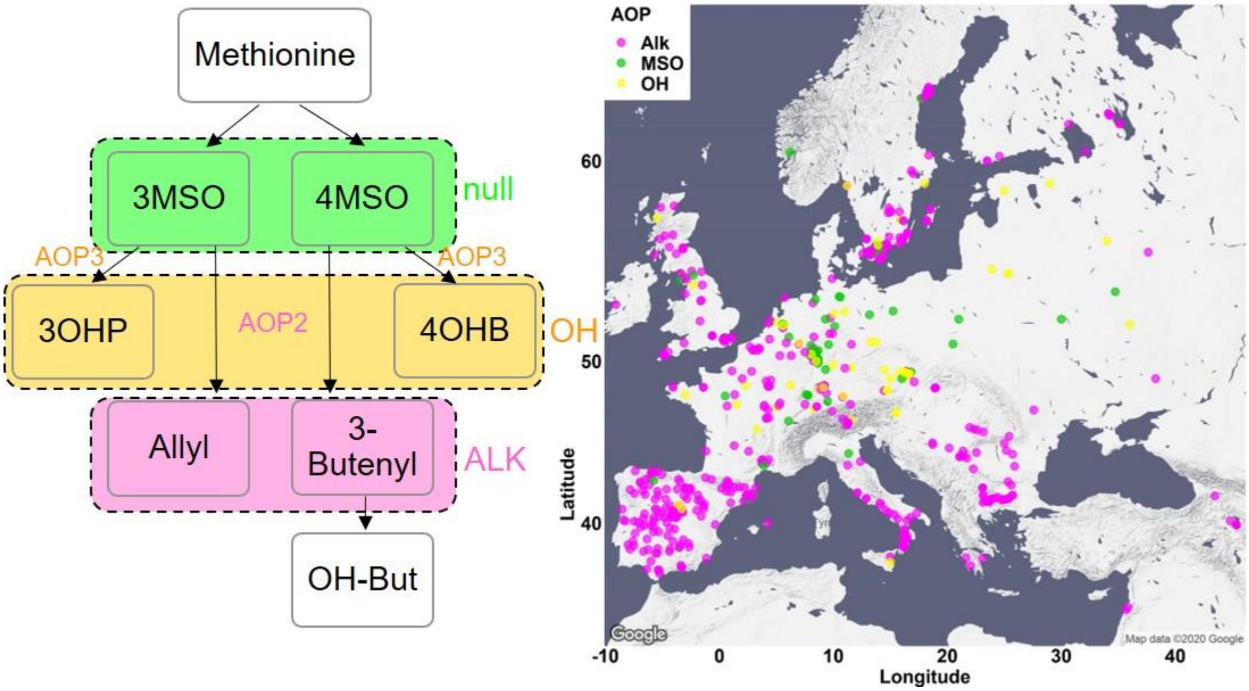


Supplementary figure 4: **Phenotypic classification based on the dominant AOP enzymes.** Relative amounts of alkenyl GSLs, alkyl GSLs and MSO GSLs were calculated in respect to the total short chained aliphatic GSLs as described in the methods section. Accessions with high amounts of alkenyl GSLs were classified as AOP2 dominant, and were colored in pink. Accessions with high amounts of alkyl GSLs were classified as AOP3 dominant, and colored in orange. Accessions with high amounts of MSO GSLs were classified as AOP null, and colored in green. The accessions were plotted on a map based on their original collection sites.

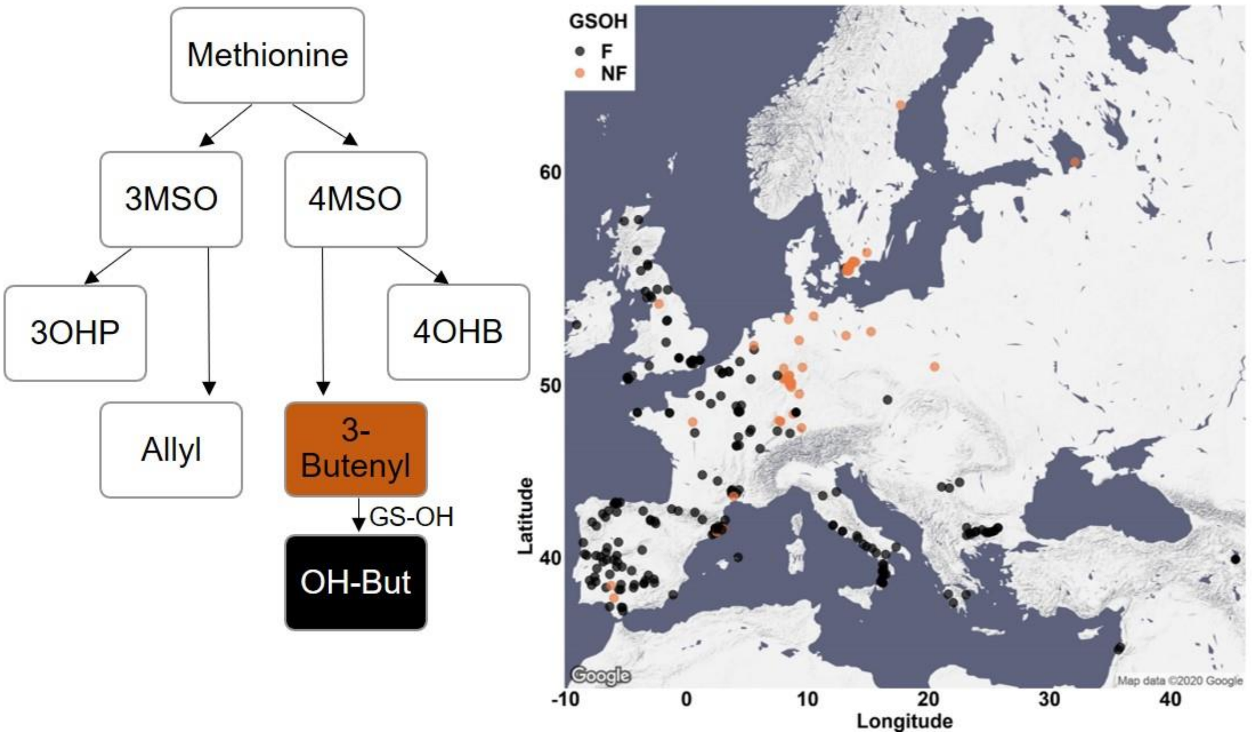


Supplementary figure 5: **Phenotypic classification based on GS-OH enzyme activity.** The ratio between 2-OH-3-Butenyl to 3-Butenyl GSL was calculated only for MAM1 dominant accessions (accessions with GSLs containing 4 carbons in their side chain). Accessions with high amounts of 2-OH-3-Butenyl were classified as GS-OH functional and colored in black. Accessions with mostly 3-Butenyl were classified as GS-OH non-functional and colored in brown. The accessions were plotted on a map based on their original collection sites.

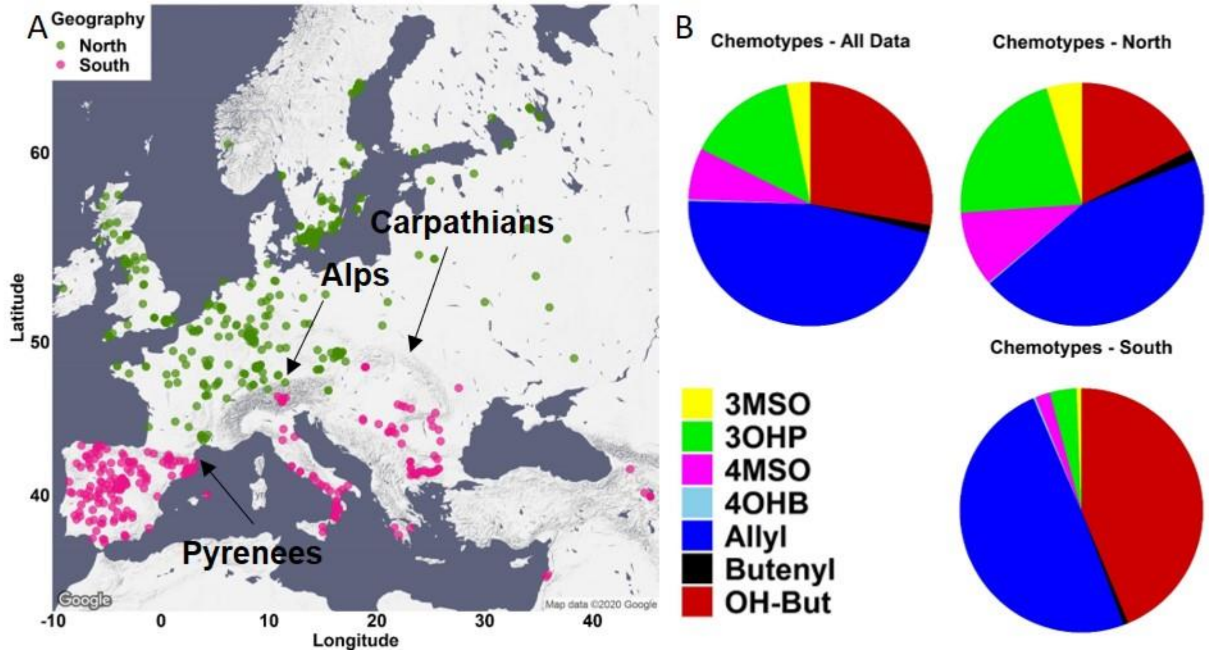


Supplementary figure 6: **Geographic partitioning of the collection.** A. The accessions were divided to two collections using the following chain of mountains: the Pyrenees between Spain and France, the Alps between Italy and Germany, and the Carpathians in the Balkan. The accessions that are located north of these mountains are referred as the northern accessions and colored in green. The accessions located south of these mountains are referred as the southern accessions and colored in pink. B. The percentage of each chemotype was independently calculated in the south and north. Butenyl = 3-Butenyl, OH-But = 2-OH-3-Butenyl.

A

| Environmental parameter | Effect on chemotype - North | Effect on chemotype - South | Interaction with Geography |
|---|---|---|---|
| Genomic Group | <0.0001 | <0.0001 | <0.0001 |
| Max temperature of warmest month | 0.0007 | <0.0001 | 0.71319 |
| Min temperature of coldest month | 0.3070 | <0.0001 | <0.0001 |
| Precipitation of wettest month | 0.9460 | 0.0012 | 0.0011 |
| Precipitation of driest month | 0.0782 | 0.6783 | 0.8711 |
| Distance to the coast | 0.5262 | 0.0022 | 0.0017 |

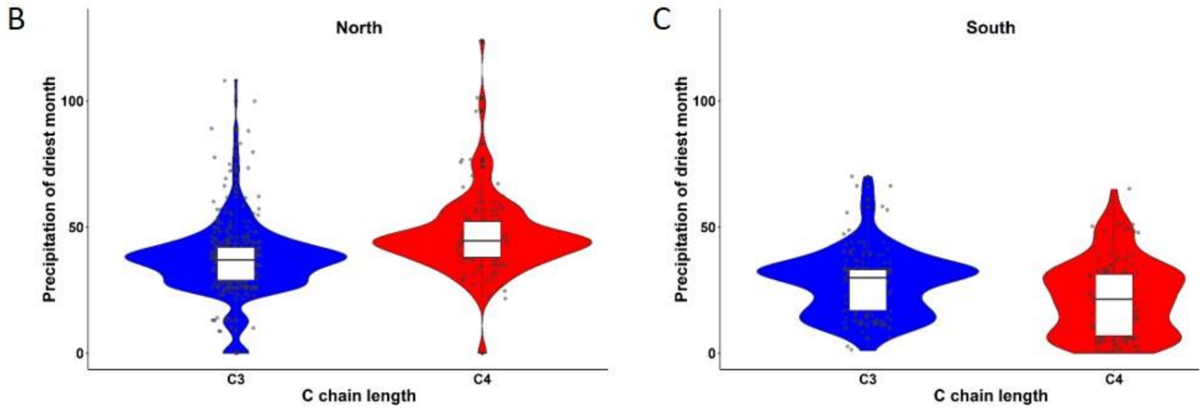


Supplementary fig 7: **Environmental conditions differentially associate with MAM status in the north versus the south.** A. Linear model for MAM status (carbon side chain length) was conducted with the indicated environmental parameters, for the northern and southern collection, separately (for more details see methods). The tables show P values for each term from the linear model. For the interaction with geography – the linear model was run using the total dataset, and the geography parameter (north or south) was added to the model. B, C. The associations of C3 and C4 accessions to precipitation values of the driest month. Significance was tested by nonparametric t-Test, P = 2.1E-09 for the North, P = 0.0006158 for the South.

| | Mean Decrease Accuracy | | | Mean Decrease Gini | | |
|---|---|---|---|---|---|---|
| | All data | North | South | All data | North | South |
| Temp. cold | 107.55 | 30.7 | 27.08 | 70.97 | 40.47 | 22.65 |
| Distance_to_the_coast | 84.22 | 38.6 | 22.64 | 62.56 | 37.75 | 19.39 |
| Genomic group | 128.35 | 69.58 | 20.77 | 61.79 | 52.59 | 13.86 |
| Temp. warm | 93.06 | 23.46 | 22.06 | 49.65 | 27.63 | 17.99 |
| Prep. Dry | 66.4 | 23.92 | 20.97 | 43.89 | 30.67 | 14.12 |
| Prec. Wet | 51.56 | 19.08 | 23.2 | 38.78 | 23.63 | 17.62 |

Supplementary fig 8: **Random Forest analyses links chemotype to environmental parameters.** Random forest analysis was performed to predict the chemotype identity based on environmental variables. The analysis was conducted with all data, just the north and just the south, separately. Mean decrease accuracy (the suitability of the variable as a predictor) and mean decrease gini (the probability of the variable to be wrong) are indicated for each one of the parameters, for each one of the datasets.

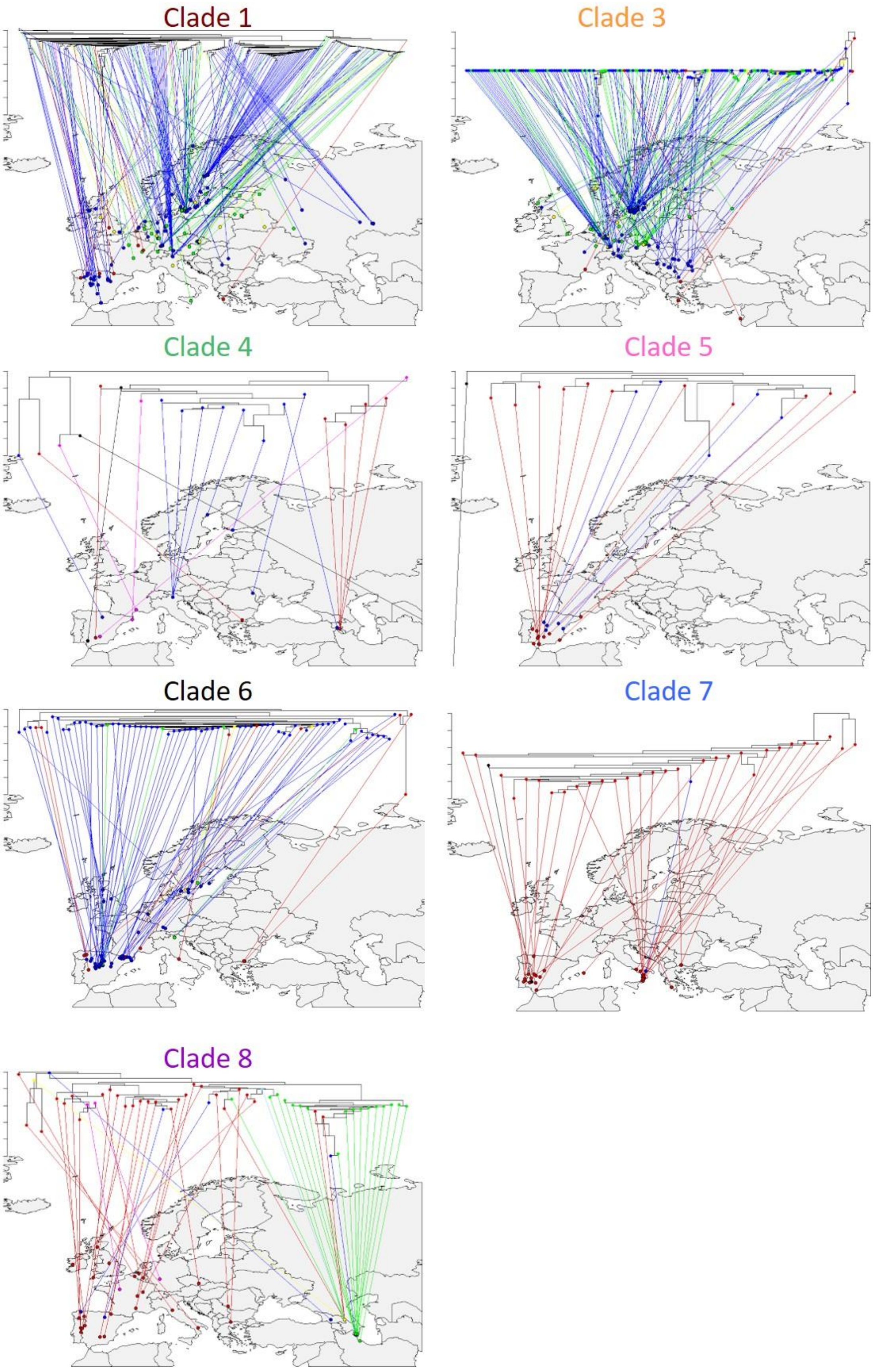
Clade 1
Clade 3
Clade 4
Clade 5
Clade 6
Clade 7
Clade 8

Supplementary figure 9: **Geographic distribution of MAM Haplotypes.** The MAM phylogeny is split by the major clades/haplotypes and each sub-clade's phylogeny is reflected on the map. Tree tips are colored based on the accessions chemotype.

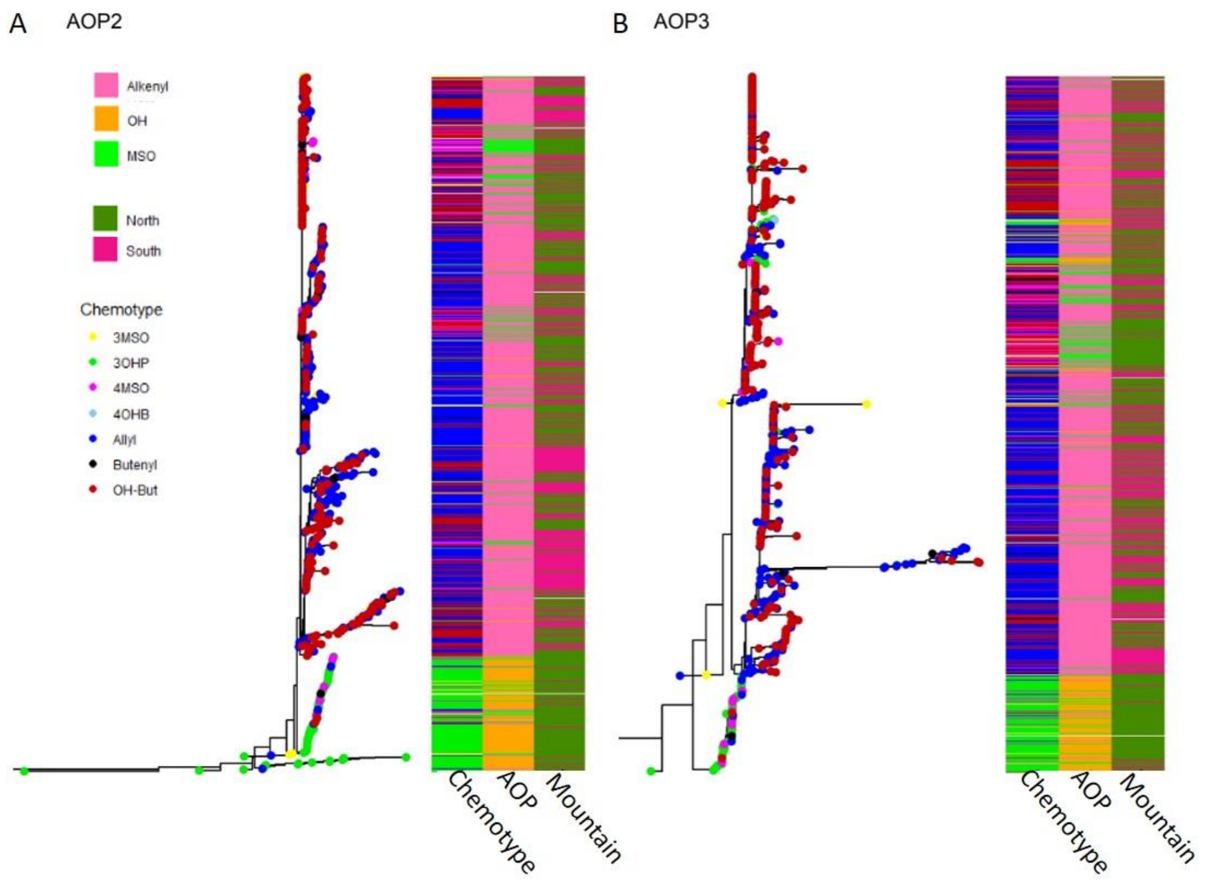


Supplementary figure 10: **AOP phylogeny.** Separate phylogenies of AOP2 (A) and AOP3 (B) across *Arabidopsis thaliana* accessions. The trees are rooted by the matching gene in *Arabidopsis lyrata, that is* not shown because of distance. Tree tips are colored based on accessions chemotype. The first column of each heatmap represents the dominant chemotype's identity. The second column of each heatmap represents the AOP functionality: pink for alkenyl (AOP2 dominant), orange for hydroxy (AOP3 dominant), green for MSO (null). The third column represent the accessions geographic location: dark green for the north, dark pink for the south.
